# A novel virus vB_PshM_Y4 with low similarity to both cultured and uncultured viruses

**DOI:** 10.1101/2025.02.10.637413

**Authors:** Zhaobin Zheng, Lei Zhao, Wei Wang, Yundan Liu, Kaiyang Zheng, Xin Chen, Suqing Zhang, Yue Sun, Zhe Ma, Hongbing Shao, Yeong Yik Sung, Wen Jye Mok, Li Lian Wong, Chen Gao, Libin Sun, Yantao Liang

## Abstract

Viruses are the abundant and diverse biological entities on the Earth. During past decades, our knowledge about genomic diversity of viruses has been greatly expanded through high-throughput sequencing. However, whether traditional isolation method could detect virus that could not be identified by high-throughput sequencing is still unknown. In this study, a novel virus, vB_PshM_Y4, infecting *Pseudoalteromonas shioyasakiensis*, which is an opportunistic pathogen of economically important marine species, was isolated. After comparing vB_PshM_Y4 with the over 15 million genomes of both cultivated and uncultivated viruses in the NCBI and IMG/VR v4 datasets, no similar genome was identified. This suggests that viral isolation methods can detect novel viruses that may not be identified through high-throughput sequencing. Transmission electron microscopy showed that vB_PshM_Y4 exhibits the myoviral morphology. The virus possesses a double-stranded DNA genome that is 158,231 bp in length, with a G+C content of 30.92%, and contains 226 open reading frames (ORFs). Six ORFs were identified as putative auxiliary metabolic genes (AMGs). Phylogenetic tree and gene content-based network analyses indicated that vB_PshM_Y4 may represent a novel viral species within an unknown family of the *Caudoviricetes* class. Additionally, biogeographical distribution analysis showed that vB_PshM_Y4 were mainly detected in the Arctic and temperate and tropical epipelagic zones with low abundance.

**Importance:** This study suggests that viral isolation could detect novel virus that could not be identified through high-throughput sequencing, contributed to a deeper understanding of the genetic diversity and distribution of *Pseudoalteromonas* viruses, and offered valuable insights into the prevention and control of opportunistic pathogenic bacteria affecting economically important marine organisms.

## Introduction

Viruses, which numerically dominate our oceans, are essential for regulating microbial populations and shaping community structures, and they have a significant impact on global biogeochemical cycles (1). Bacteriophages are viruses that infect bacteria, and studies have shown that they can kill up to 20-40% of marine prokaryotic cells each day (2). However, beyond their impact on host mortality, they also influence host diversity in various ways. They can contribute genetic material to their hosts via horizontal gene transfer (HGT) and maintain host diversity by frequency dependence (3, 4). During past decades, our knowledge about genomic diversity of viruses has been greatly expanded through high-throughput sequencing (5). Currently, the largest genomic dataset for viruses on Earth, IMG/VR v4 dataset, contained more than 15 million genome sequences about cultivated and uncultivated viruses, which is about 1000-fold of viral genomes in the International Committee on Taxonomy of Viruses (ICTV) dataset (6). Hence, one of the common sense is that high-throughput sequencing could identify more viral genomes than the traditional viral isolation methods (6). However, whether traditional isolation method could detect virus that could not be identified by high-throughput sequencing is still unknown.

*Pseudoalteromonas* is a model genus within the *Alteromonadales* order and the *Pseudoalteromonadaceae* family, consisting of rod-shaped, Gram-negative bacteria (7). They are prevalent in diverse marine environments and known for producing various bioactive compounds (8). *Pseudoalteromonas shioyasakiensis*, was first isolated from Pacific sediment in 2014 and is recognized for its ability to produce polysaccharides (9). *P. shioyasakiensis* can infect abalone and inhibit the growth of *Vibrio penaeicida*, a pathogen lethal to shrimp (10). Despite significant advancements in the study of marine viruses and the development of various experimental methods for isolating them, our understanding of the marine viruses infecting *P. shioyasakiensis* is still limited.

In this study, a novel virus, vB_PshM_Y4, infects *P. shioyasakiensis* was isolated from water samples collected in Xiaogang, Qingdao, China. The genomic and evolutionary relationships analysis of vB_PshM_Y4 showed a large variance from other cultured and uncultured viruses in the NCBI and IMG/VR v4 databases. Additionally, the characteristics and biogeographical distribution of vB_PshM_Y4 were investigated. This research contributes to the growing understanding of marine viruses and sheds light on the interactions between *P. shioyasakiensis* and viruses, offering valuable insights for the prevention and management of diseases in economically important marine species.

## Materials and methods

### Isolation and purification of host

A seawater sample was collected from Xiaogang, Qingdao, China (36.1000°N, 120.4833°E) on August 27, 2023. Subsequently, the sample was introduced onto 2216E agar medium and incubated at 25°C using the plating technique for 24 hours. Single colonies were selected and purified through at least three rounds of streak plating. After purification, the isolates were stored at 4°C (11).

### Determination of host bacteria

Determined the species of host based on 16S rRNA by the online version of EZBioCloud (https://www.ezbiocloud.net/) (12). Sequences closely related to the host were downloaded from GeneBank as references. IQ-Tree2 (v2.0.3) was used to perform a rapid bootstrap analysis with 3000 replicates to assess node support and to construct a phylogenetic tree of host bacteria with these reference sequences (13). Visualization was performed using iTOL (v5.0) (14).

### Isolation and purification of virus vB_PshM_Y4

Virus vB_PshM_Y4 was isolated using the double-layer agar method (15). Briefly, the water sample was filtered through a 0.22 μm pore size membrane to remove bacteria and planktonic algae. A 200 μL logarithmic phase culture of the host bacterium was then mixed with 200 μL of viral filtrate and incubated for 15 minutes. This mixture was combined with 4.5 mL of semi-solid agar at 45°C, poured onto the surface of a solid medium, and then cultured overnight at 26°C (16), with three replicate agar plates prepared. A plate showing clear plaques was selected, and the virus mixture was soaked in 3 mL of SM buffer (8 mM MgSO₄·7H₂O, 100 mM NaCl, 50 mM Tris-Cl, pH 7.5) at 4°C for 24 hours before being filtered through a 0.22 μm PES Millipore filter membrane. The purification process was repeated at least three times to obtain a purified virus solution, which was subsequently stored in SM buffer at 4°C.

### Observation of virus morphology

Transmission electron microscopy (TEM) was employed to classify the morphology of virus vB_PshM_Y4. In brief, 20 µL of the purified virus sample was placed on a 200-mesh copper grid and negatively stained with 1% (wt/vol) phosphotungstic acid (pH 7.0). The electron micrographs of vB_PshM_Y4 were captured at 100 kV using a TEM (JEM-1200EX, Japan) (17). The size of the virus was estimated from the electron micrographs using the built-in ruler of the TEM (18).

### One-step growth curve

To determine the latent period of vB_PshM_Y4, the double-layer agar method was employed to assess the one-step growth curve of the virus. Initially, a suspension of vB_PshM_Y4 virions was used to infect a host culture in the logarithmic growth phase at a multiplicity of infection (MOI) of 0.1, and the mixture was incubated at 28°C for 20 minutes to allow for infection. Centrifugation and washing of the culture were repeated to ensure the removal of any residual free virions, and the cells were then resuspended in 50 mL of fresh 2216E liquid medium. Samples were collected at 5-minute intervals for the first 40 minutes, at 10-minute intervals up to 60 minutes, and at 30-minute intervals up to 180 minutes. The viral titers at each time point were quantified using plaque assays(19, 20). The burst size, calculated based on the number of plaques at various time points, is quantified to reflect viral growth dynamics (21).

### Thermal stability and pH stability assay

To assess thermal stability, a virus suspension (10^8^ PFU/mL) was exposed to temperatures ranging from -20°C to 90°C for 2 hours, with data recorded at 10°C intervals. Each temperature condition was tested in triplicate, and plaque counts were recorded over four days to evaluate the impact of each temperature treatment.

To evaluate pH stability, a virus solution with an initial titer of 10⁸ PFU/mL was prepared and adjusted to pH levels ranging from 2 to 12. The solution was divided into 11 aliquots of 5 mL each, and these were cultured at 28°C for 2 hours. Following this, 200 μL of each pH-adjusted aliquot was mixed with an equal volume of host bacterial culture in the logarithmic growth phase (10⁷ CFU/mL). The mixtures were plated onto double-layer agar and incubated at 28°C for 12 hours, and then the plaque counts of each aliquot were recorded(20, 22).

### Virus DNA extraction and sequencing

The nucleic acid of vB_PshM_Y4 was extracted using the OMEGA viral DNA kit according to the instructions provided by the manufacturer, and the purified viral genomic DNA was sequenced using the Illumina NovaSeq platform with 2×150 bp paired-end sequencing, conducted by Biozeron Biotechnology Co., Ltd. in Shanghai, China.

Soapnuke (v2.0.5) was used to generate high-quality clean reads, these clean reads were aligned to the *P. shioyasakiensis* genome using BWA (v0.7.17) with default settings (mem -k30) to reduce the impact of host sequences. Any alignment results which less than 80% of the total read length matched were excluded, and the corresponding sequences were removed. Following this, the clean reads were assembled de novo using Megahit (v1.1.2) with its default parameters (--presets meta-large --min-contig-len 300).

Combined Prodigal, RAST, and GeneMarkS to predict the open reading frames (ORFs) of vB_PshM_Y4. These ORFs were then annotated through the Pfam database (https://pfam.xfam.org/search/sequence) using its default settings (23). To further refine the annotations, BLASTp searches were conducted against the nonredundant (NR) protein database (http://blast.ncbi.nlm.nih.gov) and compared the results with the KOfam database, a customized HMM database of KEGG Orthologs (16, 24). The genomic map was visualized using CLC, while the GC-skew was calculated and displayed using Genskew with a window size and step of 100 bp (25), tetranucleotide frequencies of vB_PshM_Y4 were then calculated using a step size of 5000 base pairs. Finally, tRNA sequences were identified employing tRNAscan-SE (https://tRNAscan-SE-Search-Server-(ucsc.edu)) (26). It is reported that tRNA Relative Contribution Index indicator to evaluate the potential gain of virus tRNA genes in the translation efficiency of each virus gene (27). tRCI and cosine similarity distance wered calculated, then ranked genes in vB_PshM_Y4 by these two values separately. Gene enrichment in the top-ranked bins was calculated using the hypergeo metric distribution probability and considered significant when the p-value was < 0.05 (11).

### Evolutionary status and comparative genomes analysis of vB_PshM_Y4

Amino acid-based clustering was performed by vConTACT (v2.0) (28). The NCBI dataset was used as the database to select homologous sequences with vB_PshM_Y4. To ensure accuracy, a database was created using diamond with the complete amino acid sequences of vB_PshM_Y4 and sequences from the NCBI virus database. An all-to-all BLASTp comparison was performed with a coverage threshold of 30% and an expected value threshold of 1e-10. The protein cluster mode was specified as MCL, the virus cluster mode as ClusterONE, and the overlap threshold between viral clusters was set at 0.01 for clustering analysis. The clustering results were visualized by Gephi (29), enabling to identify and select sequences with significant connections to vB_PshM_Y4 for further shared protein analysis.

To identify the phylogenetic relationships of vB_PshM_Y4, whole-genome protein tree analysis was conducted with reference sequences by ViPTree (v4.0) (genome.jp/viptree) (30). For further analysis, representative reference sequences which closely related to vB_PshM_Y4 were selected for construction of phylogenetic tree through ViPTree, along with the complete protein sequence of vB_PshM_Y4 and ViPTree reference sequences. Following that, sequences from branches closer to vB_PshM_Y4 were picked for gene collinearity analysis. All sequences and host information were obtained from the NCBI database. Virus Intergenomic Distance Calculator (VIRIDIC) was used to calculate the Average Nucleotide Identity (ANI) between the selected representative reference sequences and vB_PshM_Y4 (31).

The terminase large subunit is highly conserved throughout evolution. To further determine the evolutionary position of vB_PshM_Y4, proteins from the same cluster as the terminase large subunit in the vConTACT analysis were selected. Additionally, terminase large subunit from sequences with primary and secondary connections to vB_PshM_Y4 were included. Using a method similar to the 16S rRNA phylogenetic tree of the host bacterium for the construction of phylogenetic tree of terminase large subunit, incorporating these selected sequences.

### The distribution of vB_PshM_Y4 in the ocean

The relative abundance of vB_PshM_Y4 in marine environments was estimated using TPM (Transcripts Per Kilobase of exon model per Million mapped reads) values. This was achieved with the metagenomics tool minimap2, and the calculations were performed using CoverM (v0.3.0) with specific parameters (–min-read-percent-identity 0.95, –min-read-aligned-percent 0.75) (32, 33). The Global Ocean Viromes 2.0 (GOV 2.0) database, consisting of 154 virome databases, was divided into five viral ecological zones (VEZs), including Arctic (ARC), Antarctic (ANT), temperate and tropical epipelagic (EPI), temperate and tropical mesopelagic (MES), and bathypelagic (BATHY) (34). 24 other viruses were selected as references, all of which have a wide distribution in the ocean.

## Result and discussion

### Phylogenetic tree of 16S rRNA of host strain

16S rRNA sequences and reference sequences from the *Pseudoalteromonas* genus were used to construct a phylogenetic tree of the host bacterium. For clarity, 19 sequences closely related to the host were used in the analysis (Fig. S1). The phylogenetic tree indicated that the host of vB_PshM_Y4 is close to *P. shioyasakiensis*. The identification by EZBioCloud indicated that the host bacteria 16S rRNA has 99.23% similarity to *P. shioyasakiensis*, resulting that the host strain belongs to this species.

### Characteristics of vB_PshM_Y4

When the virus infects its host on double-layer agar, clear plaques are formed. TEM images revealed that vB_PshM_Y4 exhibits the myoviral morphology, with a head diameter of approximately 40 nm and a tail around 50 nm (Fig. 1A).

**Fig. 1.**
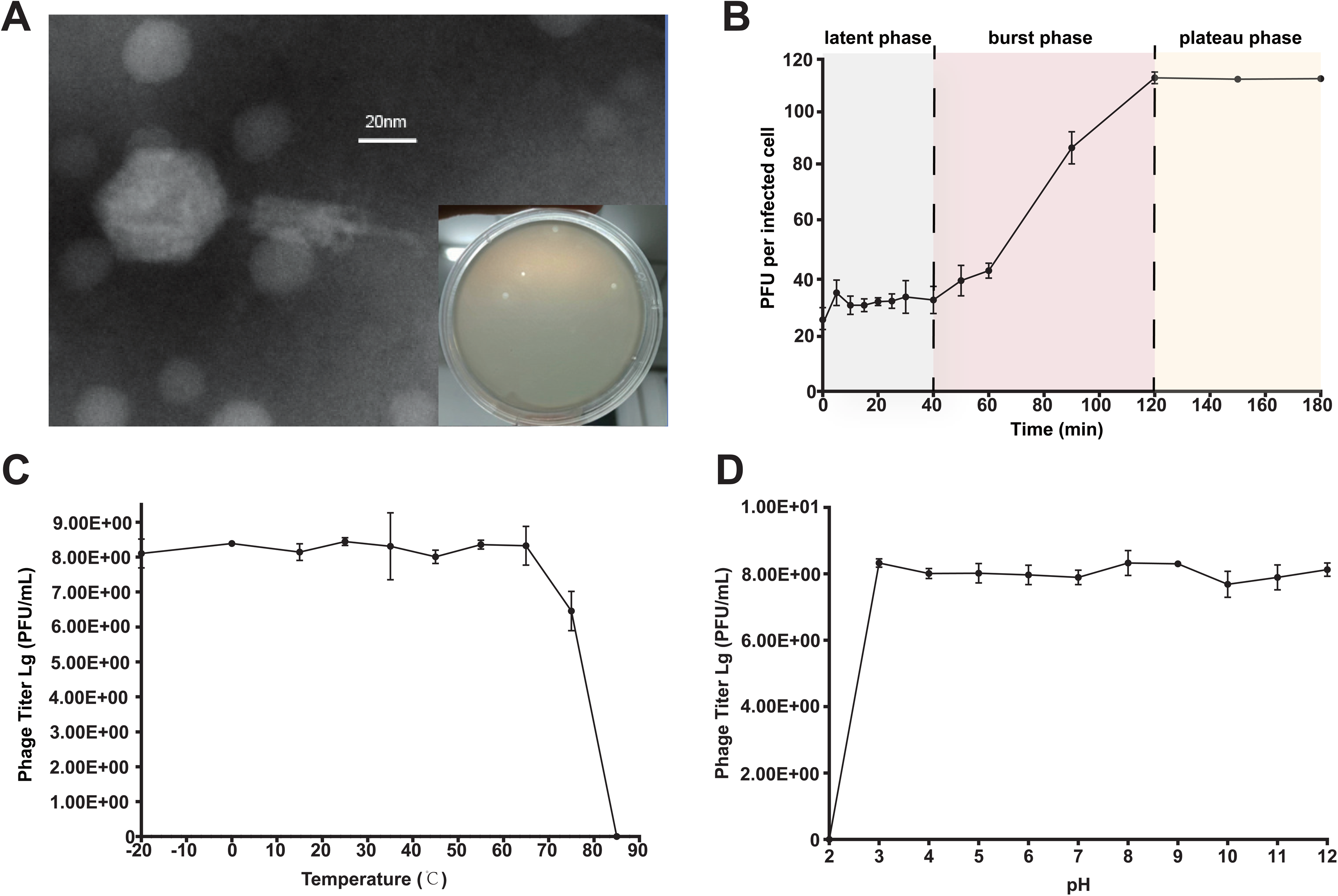
Biological properties of vB_PshM_Y4. (A) Plaque of vB_PshM_Y4 and TEM morphology of vB_PshM_Y4. Viruses were negatively stained with potassium phosphotungstate. Scale bar = 20nm. (B) One-step growth curve of virus vB_PstM_Y4. Each point represents the average value from triplicate experiments, and error bars indicate the standard deviations (SDs) of three independent experiments. (C) The curve of thermal stability (D) The curve of pH stability.

The one-step growth curve analysis of vB_PshM_Y4 indicated that the virus has a latent phase of 40 min following host infection. The burst phase occurs between 40 and 120 min, after which the viral titer stabilizes and enters the plateau phase. The average burst size is about 110 viral particles per cell (Fig. 1B).

The temperature tolerance experiments revealed that vB_PshM_Y4 reached its highest titer at 26°C. The virus maintained a stable and relatively high titer within a temperature range of -20°C to 75°C. Over 75°C, the titer declined, with the virus becoming completely inactive at 85°C (Fig. 1C). The pH stability experiments showed that vB_PshM_Y4 remained active within a pH range of 3 to 12. Below pH 3, the activity of this virus rapidly decreased, and it became completely inactive at pH 2 (Fig. 1D). The wide temperature and pH range suggested that this virus can be used to treat disease caused by *P. shioyasakiensis*.

### Genomic analysis of vB_PshM_Y4

Sequencing and assembly results indicated that the genome of vB_PshM_Y4 is a double-stranded DNA sequence consisting of 158,231 bp with a G+C content of 30.92%. A total of 14 tRNAs (Tab. 1) were identified using tRNAscan-SE. To investigate which functional gene might benefit from the presence of tRNA genes in vB_PshM_Y4, tRCI values were calculated (Fig. 2) (27). Results indicated that hypothetical genes have the highest tRCI values and benefit the most from the tRNA pool. Additionally, hypothetical genes were more abundant in the tRCI bin and enriched in all bins except bin 5, indicating that the tRNA carried by virus vB_PshM_Y4 contributes little to the translation of these genes. Similarly, using the cosine similarity distance with the host, the genes of virus vB_PshM_Y4 were divided into 10 bins. DNA replication and metabolism genes in bins 1, 2, 3, and 5 were marked as significant, while structural genes were only significant in bin1. This indicated that the expression of these genes is more reliant on the tRNA of host.

**Fig. 2.**
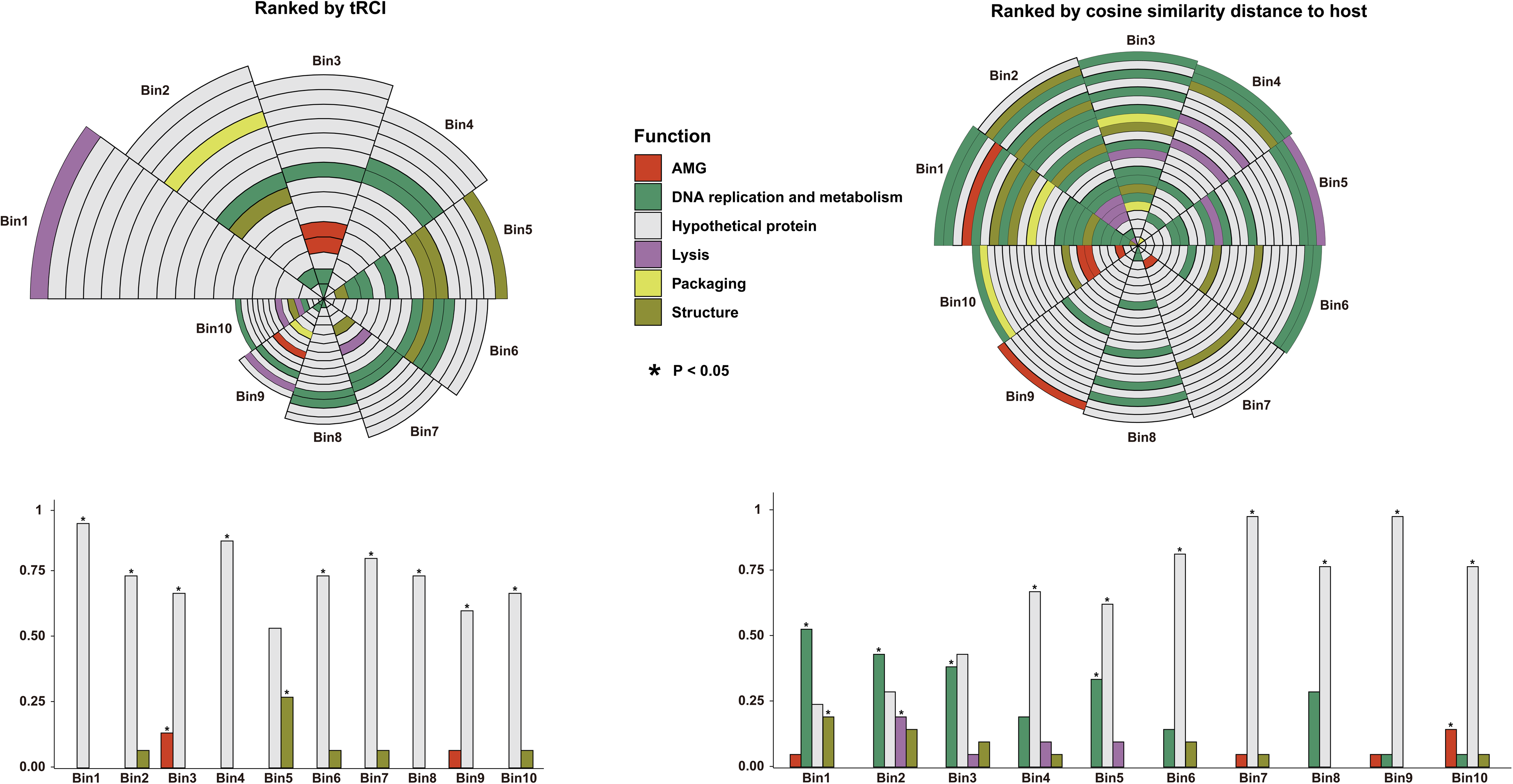
The ranks of genes with specific functions of vB_PshM_Y4 according to the cosine similarity distance to host and tRNA Relative Contribution Index (tRCI) value. Each gene in vB_PshM_Y4 is sorted according to the value of cosine similarity distance to host and assigned into 10 bins with 22 genes in each bin. Each gene in vB_PshM_Y4 is sorted according to the tRCI and assigned into 10 bins with 15 genes in each bin. A value of *p* < 0.05 indicates that the enrichment is significant.

**Table. 1.** 14 tRNAs of vB_PshM_Y4.

| tRNA id | Start | End | Type |
| --- | --- | --- | --- |
| Y4_tRNA_1 | 104814 | 104729 | Tyr |
| Y4_tRNA_2 | 104213 | 104138 | Ile2 |
| Y4_tRNA_3 | 103062 | 102989 | Pro |
| Y4_tRNA_4 | 102974 | 102897 | Lys |
| Y4_tRNA_5 | 102807 | 102732 | Ile |
| Y4_tRNA_6 | 102651 | 102577 | Glu |
| Y4_tRNA_7 | 102395 | 102319 | Asp |
| Y4_tRNA_10 | 101516 | 101442 | Asn |
| Y4_tRNA_11 | 100756 | 100686 | Trp |
| Y4_tRNA_12 | 99788 | 99715 | Cys |
| Y4_tRNA_13 | 99509 | 99436 | His |
| Y4_tRNA_14 | 99269 | 99195 | Arg |

Cumulative GC-skew analysis was performed to determine the origin and terminus of replication in the viral genome, identifying the origin at 120,800 nt and the terminus at 11,300 nt (Fig. S2). The tetranucleotide frequency analysis showed that the tetranucleotide frequencies at positions 1-10000, 25001-35000, 65001-95000, and 120001-155000 differ significantly from other locations, suggesting the presence of HGT from host at these positions.

The genome of vB_PshM_Y4 contains 226 ORFs, which have been categorized into six functional modules based on annotated protein functions: DNA replication and metabolism (n=49), structure (n=15), lysis (n=10), putative auxiliary metabolic genes (AMGs) (n=6), packaging (n=5) and unknown function (n=141). Among these ORFs, 93 are located on the positive strand, while 133 are on the negative strand (Fig. 3). The virus genome utilizes three types of start codons: ATG (n=223), TTG (n=2), GTG (n=1).

**Fig. 3.**
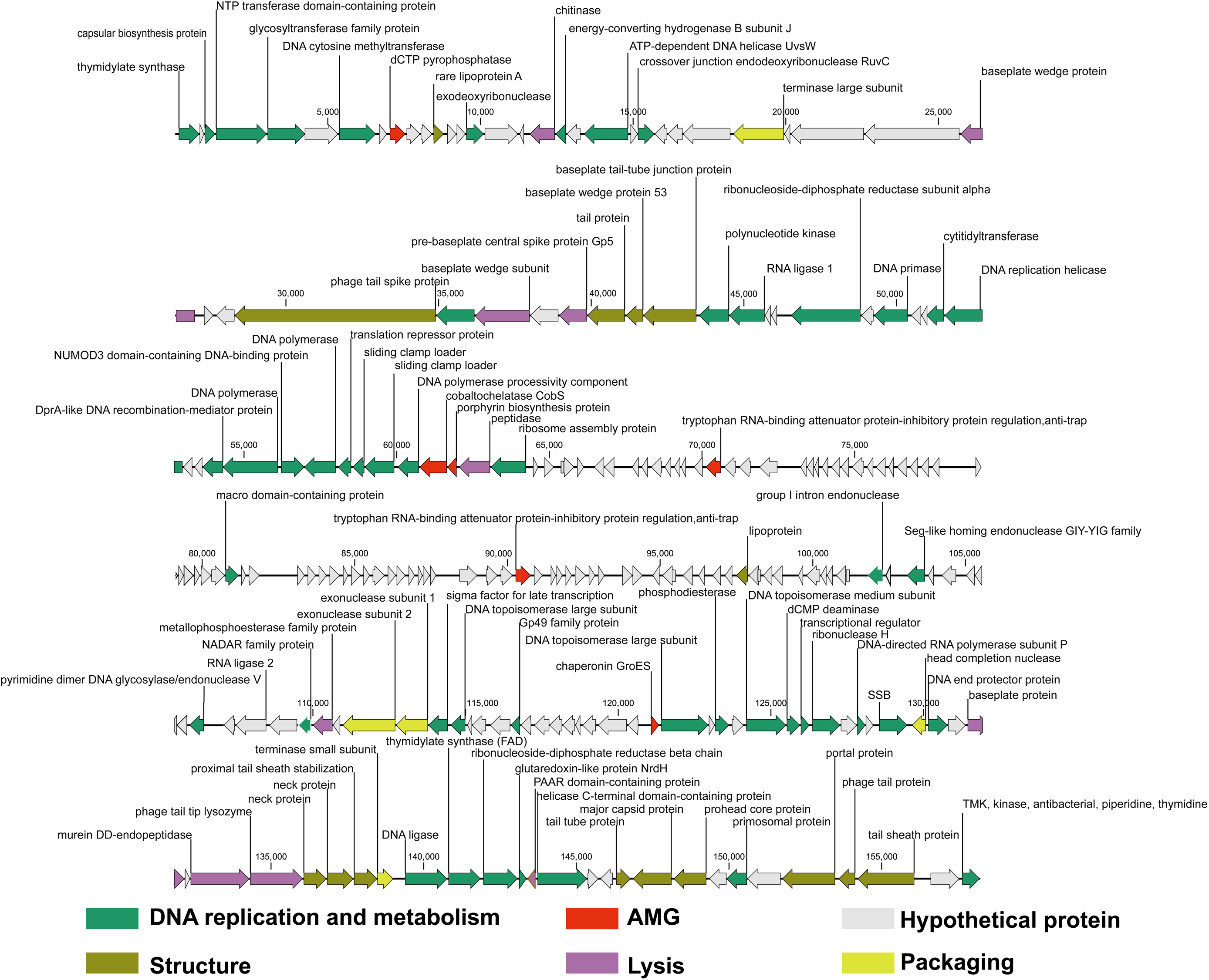
Genome map of the virus vB_PshM_Y4. Putative function categories were defined according to annotation and are represented by different colors. Each arrow represents the gene length and its transcriptional direction.

### Putative AMGs of vB_PshM_Y4

During viral infection, the expression of AMGs is thought to be involved in regulating host metabolism to enhance viral replication (35). In vB_PshM_Y4, a total of six AMGs have been annotated, encoding dCTP pyrophosphatase (ORF9), cobaltochelatase CobS (ORF63), porphyrin biosynthesis protein (ORF64), two tryptophan RNA-binding attenuator protein-inhibitory protein regulations (ORF81 and ORF125), and chaperonin GroES (ORF184).

dCTP pyrophosphatase (DCTPP1) is a phosphatase that targets deoxyribocytidine triphosphate (dCTP). It regulates pyrimidine metabolism by hydrolyzing both regular and modified dCTP. Additionally, DCTPP1 is involved in maintaining the steady-state levels of other pyrimidine nucleotides, ensuring balanced nucleotide pools within the cell (36). DCTPP1 is commonly expressed in the cytosol and nucleus of tumors, with a tendency to accumulate in the nuclei of cancer cells (37). It plays a role in restricting the concentration of non-canonical nucleotides in the nucleotide pool. The nuclear accumulation of DCTPP1 in tumor cells may be critical for maintaining proper DNA replication, thereby supporting the survival and proliferation of these cells (38).

Vitamin B12 (cobalamin) is an anti-pernicious anemia factor and one of the most structurally complex small molecules found in nature. In the oxygen-dependent (aerobic) cobalamin biosynthesis pathway, cobalt is incorporated into a tetrapyrrole structure called hydrogenobyrinic acid a,c-diamide (HBAD) by the enzyme cobalt chelatase. This enzyme is composed of three subunits—CobN, CobS, and CobT—which work together to facilitate the insertion of cobalt into HBAD, a critical step in the biosynthesis of cobalamin (39), the CobS subunit of cobalt chelatase (ORF63) has been annotated in vB_PshM_Y4. The porphyrin biosynthesis protein (ORF64) plays a role in the porphyrin synthesis pathway. Porphyrins are a class of organic compounds characterized by a tetrapyrrole ring connected by methine bridges. Their structure is similar to that of vitamin B12 and essential for the synthesis of various crucial biomolecules in organisms, including hemoglobin, chlorophyll, and vitamin B12. The presence of the CobS subunit of cobalt chelatase (ORF63) and the porphyrin biosynthesis protein (ORF64) suggested that the virus vB_PshM_Y4 may play a role in the synthesis of vitamin B12 in *P. shioyasakiensis*.

The tryptophan RNA-binding attenuator protein-inhibitory protein regulation (ORF81 and ORF125) is a protein involved in the metabolism and expression regulation of tryptophan. This protein is associated with a negative feedback regulation mechanism in the synthesis of tryptophan within cells, regulating tryptophan balance (40).

Chaperonin GroES (ORF184) is a molecular chaperone protein that belongs to the heat shock protein (HSP) family. It works together with another chaperone protein, GroEL, to form a large molecular complex. This complex helps other proteins fold correctly into their functional forms. GroES typically acts as a cap on the GroEL complex, helping to maintain an appropriate folding environment and releasing properly folded proteins when necessary. This process is crucial for cells to maintain protein stability and functionality under stressful conditions (41).

Overall, by encoding DCTPP1, vB_PshM_Y4 may create a nucleotide environment within host cells that is conducive to its own replication, while also limiting the concentration of non-canonical nucleotides to prevent interference with viral genome synthesis (42). The presence of the CobS subunit and porphyrin biosynthesis protein in vB_PshM_Y4 suggests that the virus may be involved in the synthesis of vitamin B12 within *P. shioyasakiensis* (43). Additionally, by encoding relevant proteins, vB_PshM_Y4 may optimize tryptophan metabolism within host cells, thereby promoting its own proliferation (44). Finally, through the encoding of GroES, vB_PshM_Y4 likely assists in the proper folding of proteins within host cells, ensuring the stability and functionality of key proteins during viral replication, thus enhancing the efficiency of viral replication (45).

### Virus vB_PstM_Y4 represents a novel viral species different from cultured and uncultured viruses

After comparing with the >15 million viral genomes and genome fragments in the IMG/VR v4 dataset, no similar genome was identified, suggesting vB_PshM_Y4 is totally different from uncultured viruses and viral isolation could detect novel virus that could not be identified through high-throughput sequencing. As our knowledge about genomic diversity of viruses has been greatly expanded through high-throughput sequencing technology, such as virome, metagenome and metatranscriotpmes (5). Currently, the largest genomic dataset for viruses, IMG/VR v4 dataset, contained more than 15 million genome sequences about cultivated and uncultivated viruses, which is about 1000-fold of viral genomes in the ICTV dataset (6). Hence, one of the common sense is that high-throughput sequencing could identify more viral genomes than the traditional viral isolation methods (6). In this study, result showed that traditional isolation method could detect novel virus that could not be identified by high-throughput sequencing, suggesting the significance of viral isolation to the virology, not only for viral characteristics, virus-host interactions, but also for novel viral genomes.

To determine the classification status of vB_PshM_Y4, a cluster analysis was conducted by vConTACT v2.0 with the complete protein sequence of vB_PshM_Y4 alongside reference sequences from the NCBI database. The analysis revealed that vB_PshM_Y4 did not cluster with any known virus groups, suggesting that it may represent a new virus group (Fig. 4A). To further explore the classification status of vB_PshM_Y4, a protomic tree was constructed using ViPTree. The results showed that sequences closely related to vB_PshM_Y4 belong to the *Caudoviricetes* class (Fig. S3). Consequently, vB_PshM_Y4 was classified within the *Caudoviricetes* class, although its precise family could not be determined. To refine this analysis, representative *Caudoviricetes* reference sequences from the NCBI database were selected, along with vB_PshM_Y4, to construct an additional protomic tree by ViPTree (Fig. 4B). The results indicated that vB_PshM_Y4 grouped within a major branch, with an estimated evolutionary distance of approximately 0.03. Additionally, a gene co-linearity analysis was performed with the reference sequences and vB_PshM_Y4 (Fig. 6A). This analysis showed vB_PshM_Y4 has a relatively low level of similarity with any cultured and uncultured viral genomes, which is insufficient to further classify vB_PshM_Y4 at the family level.

**Fig. 4.**
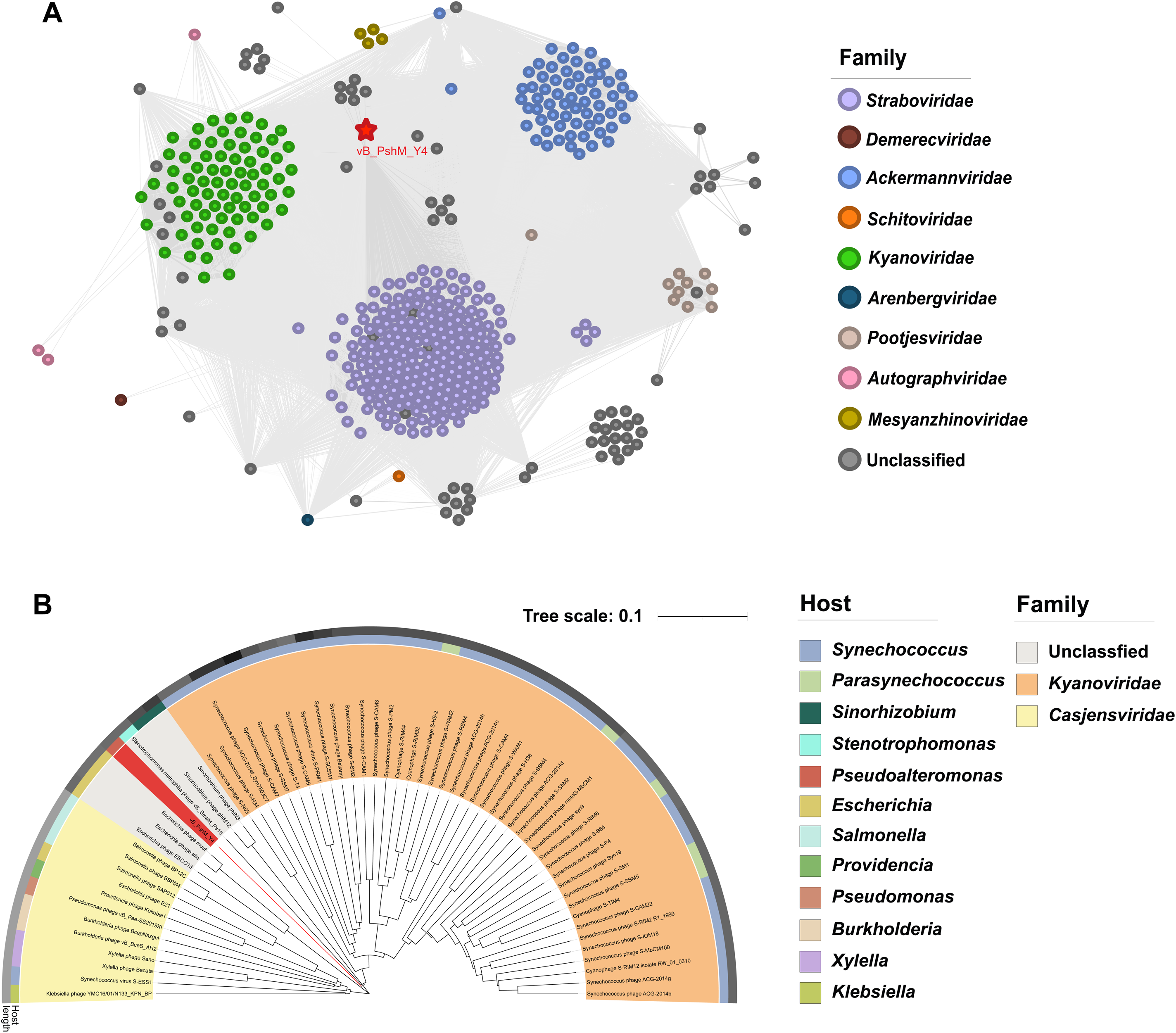
Network and protemic tree analysis of vB_PshM_Y4. (A) Protein-based viral network diagram. Viruses at different family levels are represented by different colors, with a red asterisk indicating vB_PshM_Y4. Circles enclose different virus groups. (B) Phylogenetic tree of virus vB_PshM_Y4 and the representative Caudoviricetes class reference sequences from NCBI, different colors represent different families.

**Fig. 5.**
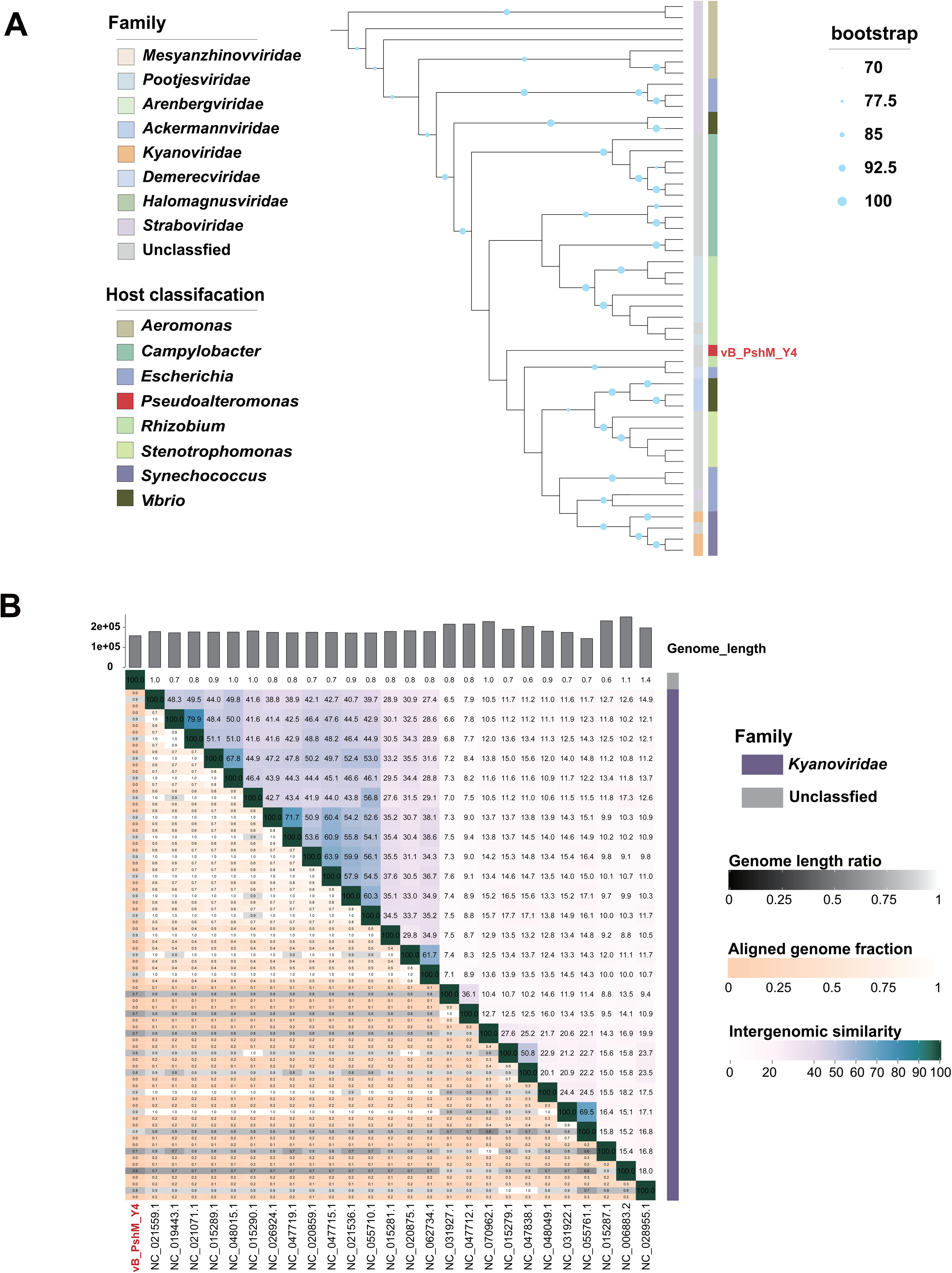
Phylogenetic and ANI analysis of vB_PshM_Y4. (A) The terminase large subunit phylogenetic tree of vB_PshM_Y4 constructed using ViPTree. For display convenience, only sequences closely related to vB_PshM_Y4 were selected. Colored ranges represent different families, and annotations in different colors represent different hosts. (B) The heatmap shows nucleotide-based intergenomic similarities of the virus vB_PshM_Y4 and reference sequences from the *Caudoviricetes* class calculated by VIRIDIC. These values represent ANI of each genome pair.

**Fig. 6.**
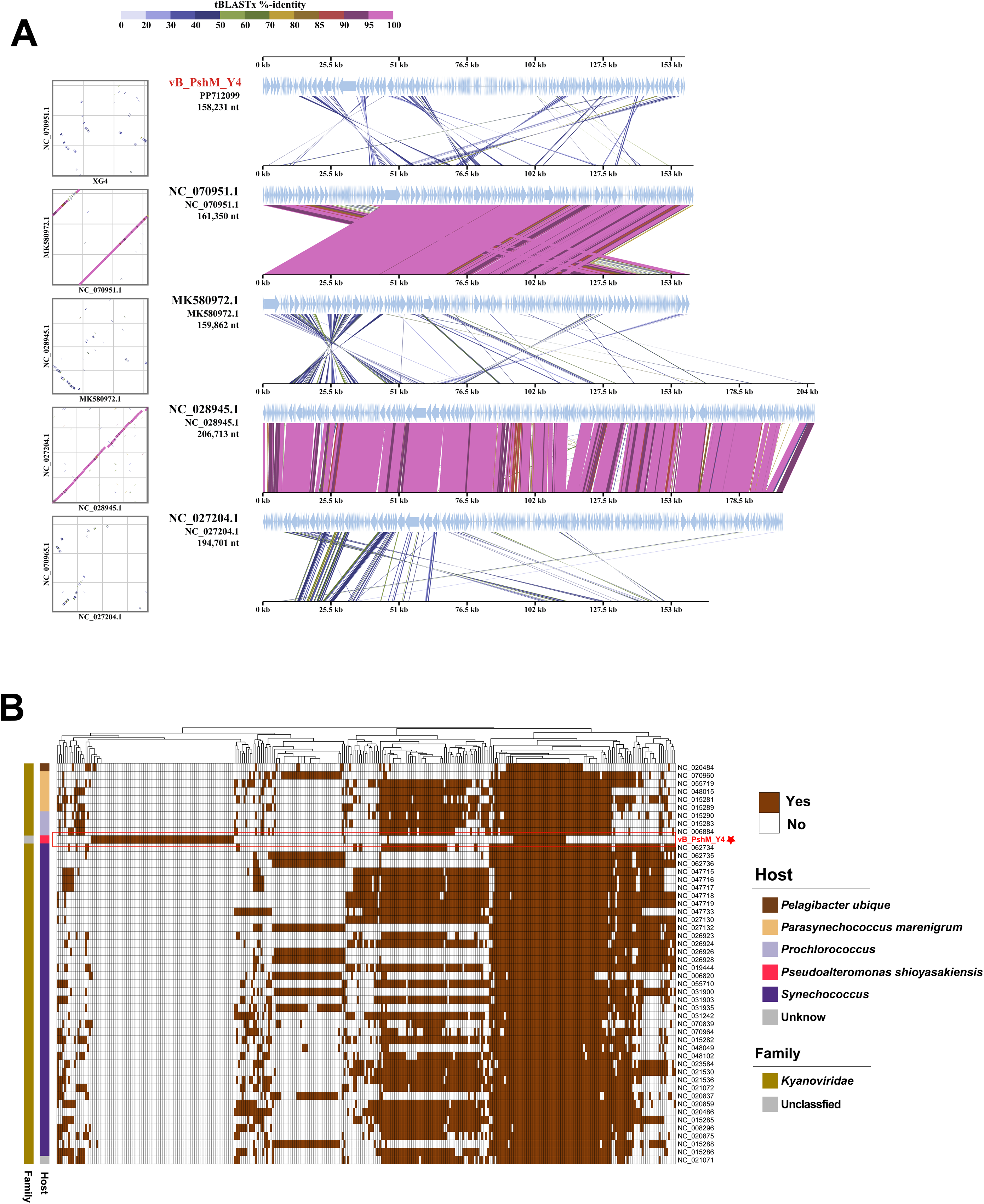
Comparison analysis of vB_PshM_Y4. (A) Genomic comparisons between virus vB_PshM_Y4 and closely related viruses in the phylogenetic tree. (B) Shared protein cluster analysis of vB_PshM_Y4 with the sequences with the highest connection to vB_PshM_Y4 from the vConTACT results. Different colors represent different families.

To further analysis the phylogeny of vB_PshM_Y4, proteins from the same protein cluster as the terminase large subunit of vB_PshM_Y4 in the vConTACT clustering results were chosen, as well as proteins with primary and secondary connections to vB_PshM_Y4, to construct the phylogenic tree of terminase large subunit. The results indicated that vB_PshM_Y4 is on a separate small branch, with a considerable evolutionary distance from the large subunit of terminal enzymes of other sequences (Fig. 5A).

In addition, VIRIDIC analysis was used to analyze the nucleotide identity of vB_PshM_Y4 and reference sequences from the *Caudoviricetes* class. The results revealed that vB_PshM_Y4 shares low similarity with any reference sequences (Fig. 5B).

Sequences with the highest connection to vB_PshM_Y4 in the vConTACT results were picked for shared protein analysis, all the selected reference sequences belonging to *Kyanoviridae* (Fig. 6B), which infect cyanobacteria (46). The results showed that vB_PshM_Y4 shares low percent of its proteins with other viruses in the *Kyanoviridae* family. And because of the low ANI between other sequences, vB_PshM_Y4 cannot be assigned to any existing family. Here, we proposed a new viral species under *Caudoviricetes*, vB_PshM_Y4, which might represent a novel viral family.

### Biogeographical distribution of vB_PshM_Y4 in the ocean

To explore the biogeographical distribution of vB_PshM_Y4 in the ocean, a feature description was conducted using 154 viral metagenomes from the Global Ocean Virome 2.0 (GOV2.0) database (34). *Pseudoalteromonas* is one of the most abundant marine bacterial genera, widely distributed in deep-sea environments, polar regions, and various marine sediments, and capable of producing a variety of bioactive compounds (47, 48). The results revealed that vB_PshM_Y4 is distributed in both the Arctic (ARC) and temperate and tropical epipelagic (EPI) zones, while its abundance is zero in the Antarctic (ANT), temperate and tropical mesopelagic (MES), and bathypelagic (BATHY) zones (Fig. S4).

## Conclusion

In this work, a unique virus vB_PshM_Y4, infecting *P. shioyasakiensis*, have been isolated and classified into *Caudoviricetes* class through genomic analysis. The relatively broad pH stability and temperature stability indicated that vB_PshM_Y4 can adapt to extreme environments and has the potential to control pathogenic bacteria affecting economically important marine animals. The six AMGs present in vB_PshM_Y4 indicated that it can influence the metabolism of host through HGT. Phylogenetic and network analysis suggest vB_PshM_Y4 is different from any cultured and uncultured viruses, and represent a novel viral species of an unknown viral family within *Caudoviricetes* class. A BLASTn analysis was performed between 56,289 vOTUs from all recorded isolates and 7,200,759 vOTUs from UViGs. The results showed that 10,450 vOTUs were present in both databases, indicating that approximately 18.56% of the vOTUs from the isolates were detected through bioinformatics methods, while only 0.15% of the vOTUs from UViGs were isolated. This result demonstrates that both bioinformatics analysis and experimental isolation are indispensable for virus research, suggesting viral isolation could detect novel virus that could not be identified through high-throughput sequencing. In conclusion, isolation and characterization of virus vB_PshM_Y4 contributes to the study of marine viruses and helps better understand the complex interactions between *P. shioyasakiensis* and virus in the ocean.

## Data availability

The annotation results and pertinent details were submitted to GenBank for the procurement of an accession number. The comprehensive genome sequence of virus vB_PshM_Y4 can now be accessed within the GenBank database, assigned the accession number PP712099.

## Authors Contributions

YL (Yantao Liang) planned, supervised, and coordinated the study and revised the manuscript. ZZ, LZ and WW performed virus isolation, performed main experiments, and bioinformatic analyses, annotated the genome, and drafted the manuscript. ZM, XC and SZ take TEM figures of virus vB_PunP_Y3. YS, KZ and YL (Yundan Liu) conducted the data curation and visualization. YL (Yundan Liu), LS, CG and HS critically evaluated the manuscript.

## Acknowledgments

We extend our gratitude to the Center for High-Performance Computing and System Simulation at the Pilot National Laboratory for Marine Science and Technology (Qingdao) for their support with high-performance server resources. We also acknowledge the support of the High-Performance Biological Supercomputing Center at the Ocean University of China, the computational resources provided by IEMB-1, a high-performance computing cluster managed by the Institute of Evolution and Marine Biodiversity, as well as the Marine Big Data Center at the Institute for Advanced Ocean Study, Ocean University of China.

## Funding information

This work was supported by the Laoshan Laboratory (No. LSKJ202203201), Natural Science Foundation of China (No. 42120104006, 42176111, and 42306111), the Ocean Negative Carbon Emissions (ONCE), and the Fundamental Research Funds for the Central Universities (No. 202172002, 201812002, 202072001 and Andrew McMinn).

## Conflict of Interest

The authors declare that they have no conflict of interest regarding this study.

## Ethical Approval

This article does not contain any studies with animals or human participants performed by any of the authors.

